# *In vitro* evolution predicts emerging CoV-2 mutations with high affinity for ACE2 and cross-species binding

**DOI:** 10.1101/2021.12.23.473975

**Authors:** Neil Bate, Christos G Savva, Peter CE Moody, Edward A Brown, Jonathan K Ball, John WR Schwabe, Julian E Sale, Nicholas PJ Brindle

## Abstract

Emerging SARS CoV-2 variants are creating major challenges in the ongoing Covid-19 pandemic. Predicting CoV-2 mutations that increase transmissibility or immune evasion would be extremely valuable in development of broad-acting therapeutics and vaccines and prioritising viral monitoring and containment. Using *in vitro* evolution, we identify a double mutation in CoV-2 receptor binding domain (RBD) that increases affinity for ACE2 almost 20-fold. We determine the mutant:ACE2 structure to reveal the binding mechanism and show the main affinity driver, Q498H, boosts binding of other RBD variants. We find this mutation incompatible with the common N501Y mutation, but N501Y variants can acquire Q498R to access a similar bonding network and affinity gain. We show Q498H, and Q498R plus N501Y, enable variants to bind rat ACE2 with high affinity. These mutations are now emerging in CoV-2 variants, such as the Omicron variant, where they would be expected to drive increased human-to-human and cross-species transmission.

## Introduction

Covid19 emerged in late 2019 and has led to more than 270 million cases of infection worldwide as of December 2021^1^. The disease is caused by the betacoronavirus severe acute respiratory syndrome coronavirus-2 (SARS CoV-2) and arose by zoonotic transmission of the virus from an animal reservoir, most likely horseshoe bat (*Rhinolophus*) via an intermediate host^2^. Since entering the human population, the virus has continued to evolve and adapt to maximise its fitness. This is leading to the emergence of viral variants with increased transmissibility as well as the ability to evade therapeutics and vaccine-induced antibodies^3^. Many of the mutations found in these variants are localised to the viral spike protein, which binds to the host cell receptor angiotensin converting enzyme-2 (ACE2) enabling viral infection^4,5^.

In addition to modifying viral transmissibility and immune escape, mutations in the spike protein can also enable the virus to bind ACE2 in species not previously susceptible to infection. Extending the range of species that a virus can infect allows the emergence of new host species and can have important health implications by creating new viral reservoirs with the potential to re-infect the human population. For example, CoV-2 has been transmitted to mink and spilled back from mink to humans^6^. This potential for cross-species transmission is especially pertinent when viruses infect species that live in close proximity to humans. Importantly, viral evolution in these new reservoirs, and recombination with other coronaviruses, has the potential to create further novel variants, that could potentially cross back into humans.

As the number of SARS-CoV-2 infected individuals rise, in human and other hosts, the opportunities for the appearance of viral mutations resulting in enhanced infectivity or pathogenesis increase. With rising immunity to the virus, there is growing selective pressure on the virus for immune evasion. Consequently, viral variants are emerging with spike protein mutations that enable antibody escape, such as the K417N mutation^3^. Some immune escape mutations lead to a decrease in binding affinity of spike protein for ACE2, and for such mutations, successful viral lineages are those that have combined the escape mutation with additional mutations that restore binding affinity. An example of this is the CoV-2 B.1.351 (Beta) variant that contains K417N and N501Y substitutions. K417N suppresses binding of a number of antibodies^3^, but also decreases binding affinity to ACE2 by around five-fold^7,8^. The N501Y substitution enhances binding affinity and can restore the decrease in affinity caused by K417N^7,8^. With selective pressure for immune escape, mutations that increase binding affinity of the spike protein become more important as they enable the virus to explore a wider range of escape mutations, including those that would otherwise hamper binding to ACE2.

The ability to predict the combinations of mutations that can arise in SARS CoV-2, and understand how these affect transmissibility and other functions, is critical for development of broadly acting therapeutics and vaccines that will be effective as the virus evolves. This knowledge is also crucial for early identification of variants that should be prioritised for monitoring and containment. Deep mutational scanning of the receptor binding domain (RBD) of SARS CoV-2 spike protein has provided valuable insights into potential effects of individual mutations affecting stability and affinity for ACE2^9^. *In vitro* evolution is another powerful methodology that can be used to identify mutations that affect protein activities, and it is particularly effective for revealing combinations of mutations that work together to modify functions.

We have been using a cell surface display evolution approach to predict potential new variants of spike protein RBD with functional effects of concern. With this approach we identify an RBD mutant containing S477N and Q498H substitutions that exhibits an almost 20-fold increase in binding affinity for human ACE2. We solve the mutant RBD:ACE2 structure to reveal the binding mechanism. Q498H dominates the affinity gain of the double mutant, and we show this mutation also boosts binding of other RBD variants. Surprisingly however, Q498H inhibits RBD binding if a CoV-2 variant already has the N501Y mutation, due to a clash between the aromatic side chains. In contrast, the related Q498R mutation is compatible with N501Y, resulting in a similar intermolecular bonding pattern as Q498H alone, and corresponding affinity gain. Importantly, we find that Q498R plus N501Y, as well as Q498H, also enable variants to bind rat ACE2 with high affinity, opening CoV-2 transmission routes for such variants between humans and rodents. Our data show Q498 to be a pivotal position by which CoV-2 can access large affinity gains and cross-species binding via two alternative mutational routes, with route selection determined by whether the variant already has the N501Y mutation. These mutations have now been found in new CoV-2 variants, such as the recently described B.1.1.529 (Omicron) variant, and are likely to drive increased variant transmissibility in the human population and potentially facilitate cross-species transmission.

## Results

The DT40 cell surface display system^10,11^ was used to identify potential spike protein variants with increased affinity for ACE2. To do this DT40 cells were transfected with DNA encoding a fusion protein consisting of the secretory leader sequence from CD5, the SARS CoV-2 RBD (amino acids 319-541) a linker region and a FLAG-epitope tag followed by the transmembrane domain and short intracellular fragment (residues 514-562) of platelet-derived growth factor receptor-β (Fig. 1a). Transfected cells were selected for antibiotic resistance and clonal populations that were positive for incorporation of the RBD construct in the rearranged immunoglobulin light chain genomic locus selected following PCR, as described previously^10^. ACE2 binding to cell surface RBD was determined by incubating cells with the indicated concentrations of monomeric ACE2 (residues 19-615, with a HA epitope tag and C-terminal Histidine_6_) and bound ACE2 detected with anti-Histidine_6_, along with RBD expression level detected using anti-FLAG. Pools of cells with the highest ACE2 binding were selected by taking diagonal sort windows to normalize for RBD expression (Fig. 1b). The number of rounds of selection was limited to three to favour selection of variants with a minimal number of accessible mutations, and sort windows were set to capture variants with maximum affinity gain. DNA encoding RBD was amplified by PCR from genomic DNA of the selected cells and sequenced. The dominant variant recovered was designated RBD4.8 and had the double substitutions S477N and Q498H (Fig. 1c). These mutations are located at each end of the binding interface with ACE2 (Fig. 1d).

**Figure 1.**
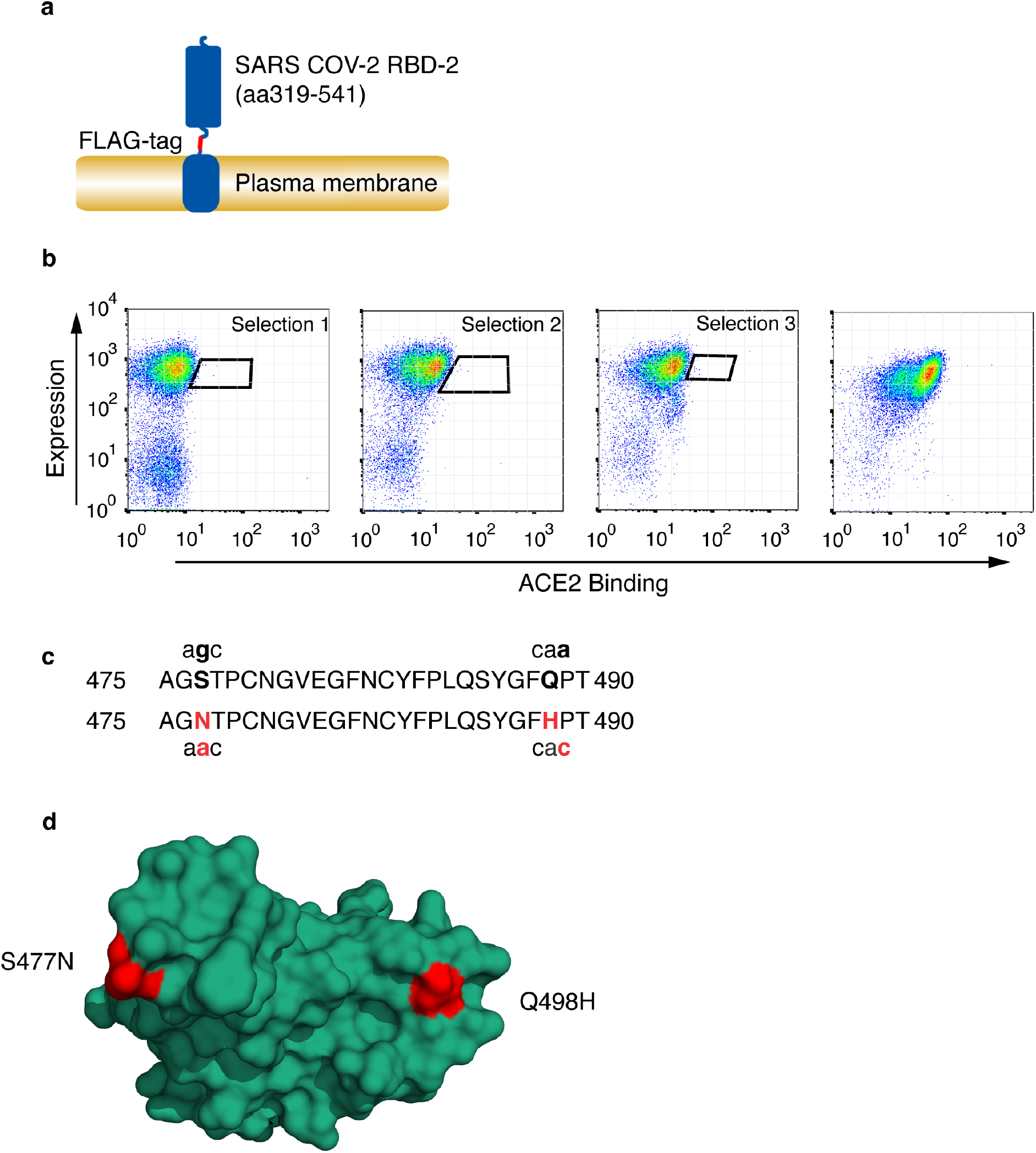
Selection of CoV-2 receptor binding domain with high affinity for HuACE2. **(a)** Schematic representation of cell surface expressed CoV-2 RBD showing RBD, FLAG-epitope tag and PDGF-receptor transmembrane anchor. **(b)** CoV-2 RBD was selected for enhanced ACE2 binding by somatic hypermutation and cell surface display in DT40 cells. FACS plots are shown of DT40 cells following binding of H_6_-tagged-HuACE2. Bound ACE2 was detected with anti-Histidine_6_-PE and expression level of RBD was assessed with ant-FLAG-APC. Sort windows are indicated for each round of selection and diversification, selecting at each round for highest Ang2 binding. Selections were performed at 0.1nM ACE2. The final panel shows a binding plot of the selected cells. **(c)** Amino acid substitutions S477N and Q498H in the selected RBD are shown, along with the corresponding single nucleotide mutations. Substituted residues and nucleotides are in red. **(d)** Position of residue substitutions in the selected RBD. Substituted residues are shown in red on the ACE2 binding interface (PDB accession number 6M0J, ^20^). The RBD is orientated to show the face that interacts directly with ACE2.

S477N and Q498H mutations have already been reported in isolates from patients. As of December 2021, S477N containing SARS-CoV-2 variants have been found in multiple countries and this mutation has emerged at least seven times during the pandemic^12^. The Q498H mutation is currently much rarer, with 37 cases being reported as of December 2021, and around 4000 cases, and rising sharply, with the related Q498R substitution, most of these being in the Omicron variant which also has the S477N mutation^12^. Infections have occurred worldwide. The co-occurrence of S477N and Q498H has yet to be reported.

### SARS CoV-2 RBDs with Q498H substitutions have high affinity for ACE2

The binding ability of soluble epitope-tagged RBD-Wuhan-Hu-1 (RBD1) and mutant RBD (Fig. 2a) was measured using biolayer interferometry (BLI) with human ACE2 (Hu-ACE2) immobilized on the biosensor (Fig. 2b). RBD1 binds Hu-ACE2 with a K_D_ of 21.2nM, consistent with previous reports^7,13,14^, whereas RBD4.8 bound Hu-ACE2 with a K_D_ of 1.1nM, representing around 19-fold higher affinity of binding than RBD1 (Fig. 2b, c). To assess the contribution of each substitution to the overall increase in affinity, we assayed the Q498H and S477N substitutions individually. RBD with Q498H bound ACE2 with a K_D_ of 2.0nM. This affinity gain of more than ten-fold is one of the largest increases in affinity seen with a single substitution in CoV-2 RBD, and this dominates the increased affinity of RBD4.8. The S477N mutant was found to bind ACE2 with a K_D_ of 4.3nM (Fig. 2b, c). In RBD4.8, N477 and H498 are positioned at each end of the RBD binding interface and work together to stabilize the RBD:ACE2 complex, once formed, as indicated by the more than nine-fold decrease in K_off_ for the combined mutant (Fig 2b,c).

**Figure 2.**
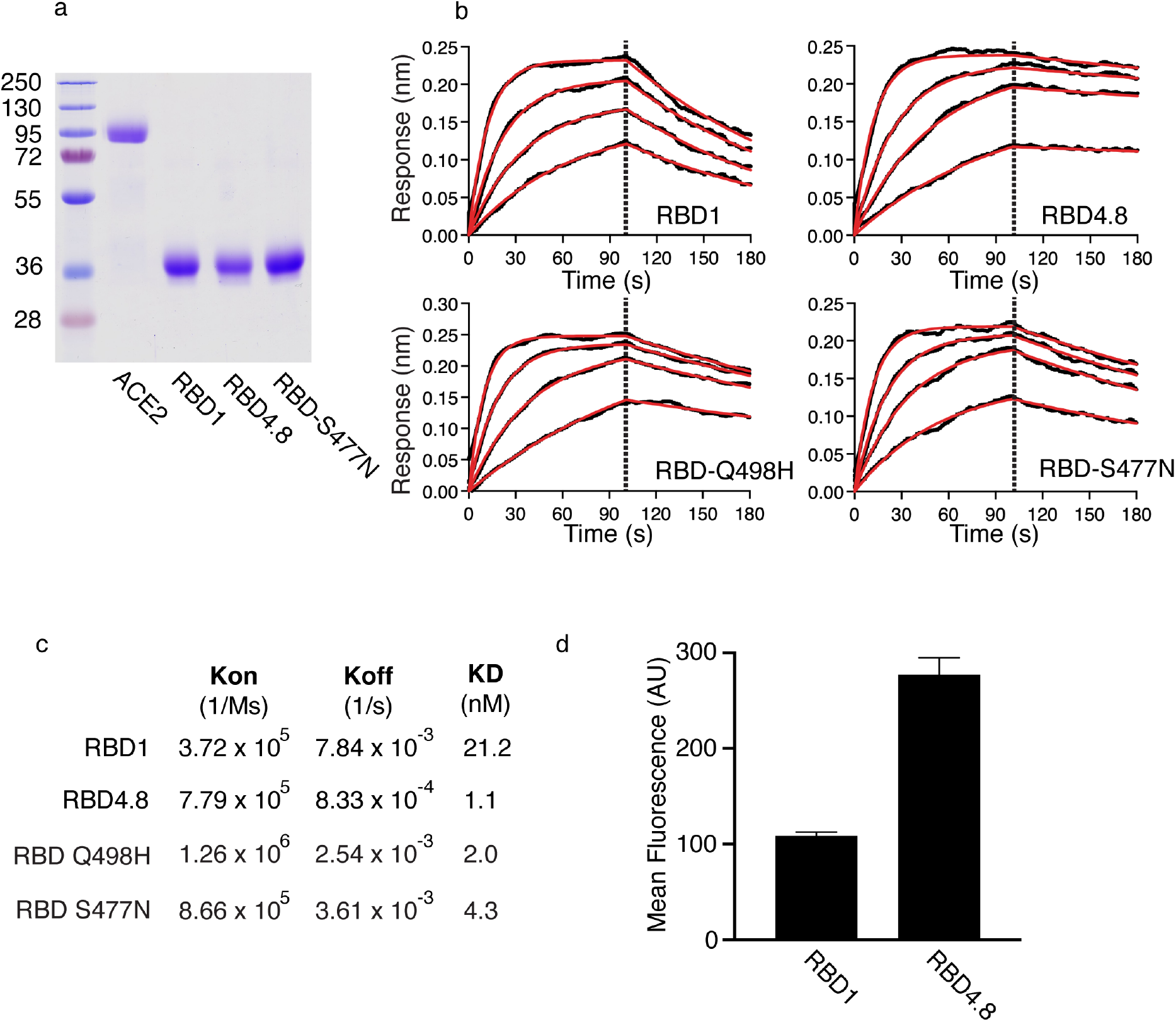
Binding affinity of CoV-2 receptor binding domain for HuACE2. **(a)** Soluble forms of HuACE2, RBD1 and mutant RBD were expressed in HEK 293 cells and purified on nickel affinity columns. Purified proteins were resolved by SDS/PAGE and detected by Coomassie staining. Molecular masses are shown in kDa. **(b)** Biolayer interferometry plots of kinetics of binding RBD to HuACE2. RBD1, or the indicated RBD, was in solution and Hu-ACE2 immobilized on sensors. Fitted curves are in red. **(c)** Kinetic binding constants for RBD binding to HuACE2 measured using biolayer interferometry. **(d)** Cellular binding of RBD and RBD4.8. Vero-E6 cells were incubated with 5μg/ml RBD or RBD4.8 at 37°C for 15 mins before washing, antibody staining of bound RBD with fluorescently conjugated antibody, and flow cytometry. Data is shown as mean fluorescence intensity and SD for three determinations.

To test whether the increase in binding of RBD4.8 for ACE2 seen in BLI assays is reflected in an increase in binding in the cellular context we used flow cytometry (Fig. 2d). Vero-E6 cells, which express ACE2, were incubated with RBD1 or RBD4.8, cells were washed and bound RBD was detected with a fluorescent antibody recognizing the epitope tag on the RBD proteins. The mean fluorescence intensity of the cells was measured by flow cytometry. Consistent with the *in vitro* binding to purified ACE2, RBD4.8 bound at increased levels compared with RBD1 (Fig. 2d).

### Mechanism of enhanced binding to Hu-ACE2

The increase in affinity of S477N/Q498H substituted RBD, and particularly the substantial increase in binding due to the Q498H substitution, prompted us to determine the structure of the RBD4.8:ACE2 complex to gain insight into the binding mechanism. The complex was imaged by Cryo-EM to produce a good quality 3.2Å map for which we could build an atomic model for the complex and observe the new interactions across the binding interface (Fig. 3a,b).

**Figure 3.**
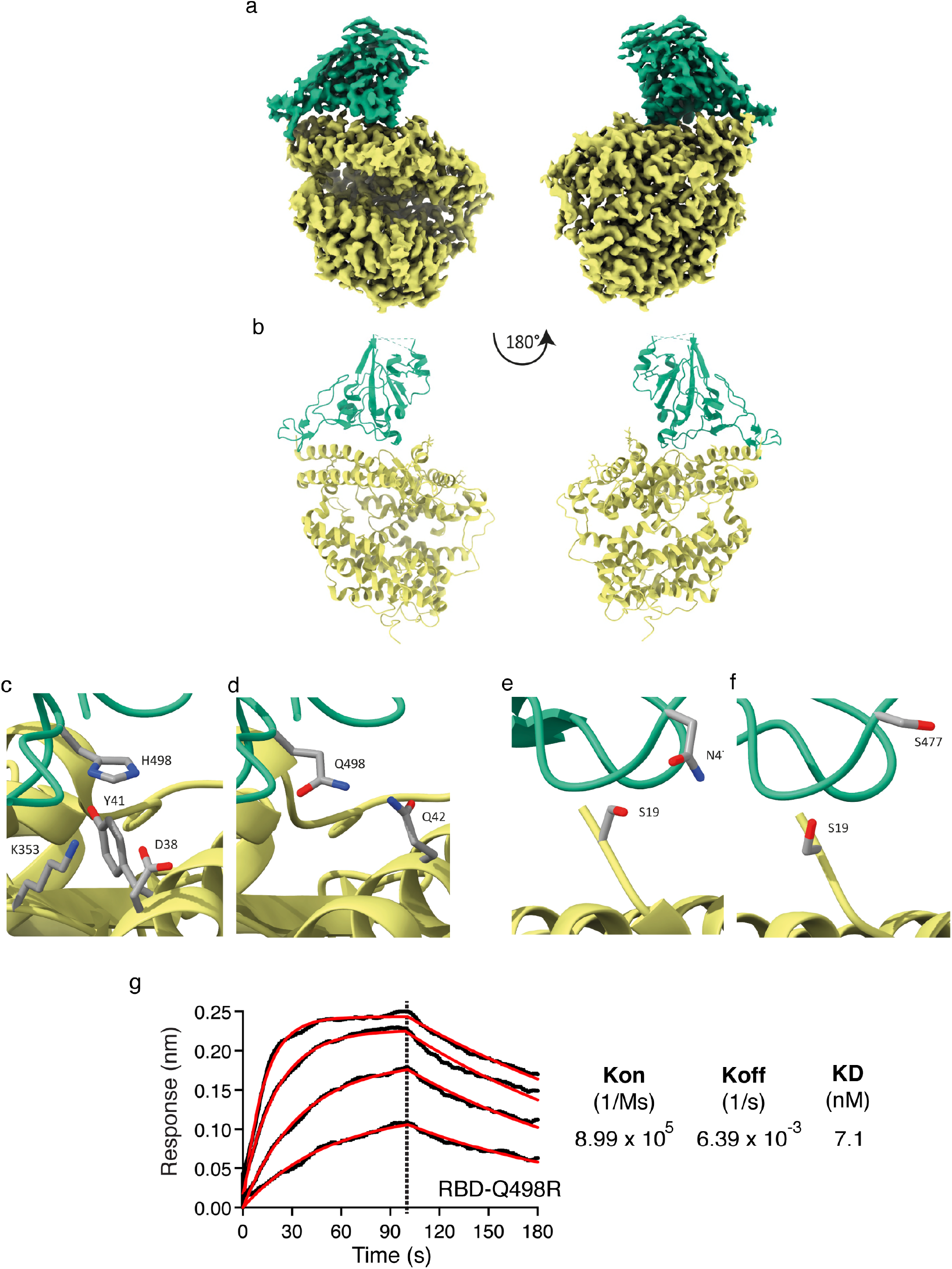
Structure of RBD4.8 in complex with HuACE2. Cryo-EM structure and model of the ACE2-RBD complex. Sharpened Cryo-EM map **(a)** of the HuACE2-RBD4.8 complex with ACE2 coloured yellow and RBD in green. The refined coordinates shown in cartoon representation **(b)** and coloured as above. In panel **(c)** H498 can be seen in proximity to ACE2 Y41 forming a non-planar π-interaction while ACE2 residues K353 and D38 are within hydrogen bonding distance to H498 and could contribute to the tighter interaction formed by this RBD mutant, whereas Panel **(d)** shows Q498 in RBD1 (PDB accession number 6M0J, ^19^) in proximity to ACE-2 Q42. Panel **(e)** indicates that the S477N substitution places this longer side-chain closer to S19 in ACE2 and within hydrogen bonding distance thus enhancing the binding between HuACE2 and RBD4.8. Panel **(f)** shows positioning of S477 in RBD1 and ACE2 S19 (PDB accession number 6M0J^19^). **(g)** Biolayer interferometry was performed with RBD containing the Q498R substitution in solution and HuACE2 immobilise on the sensor. Curves were fitted and used to calculate K_on_, K_off_ and K_D_.

Examination of the structure reveals substitution of RBD Q498 for histidine results in a perpendicular Y-shaped π-interaction between ACE2 Y41 and RBD H498^15^ (Fig 3c). In addition, rbdH498 now lies within hydrogen bonding distance of a flexible ACE2 lysine residue (K353) which would strengthen the interaction in this area. A neighbouring aspartate side chain (ACE2 D38) can also mediate a hydrogen bond via a water molecule to RBD H498 and this is supported by a continuous density between ACE2 D38 and RBD H498 at lower contour levels. Though we have not built water molecules at this resolution, several densities in our map coincide exactly with the water molecules built for the x-ray model used as starting coordinates for this study (PDB 6M0J). On the other side of the interface between ACE2 and RBD the gain of a carbon atom from the RBD S477N mutation places this side chain in hydrogen bonding distance of ACE2 S19 (Fig.3e), whereas S477 in the un-mutated RBD is unable to reach S19 (Fig. 3f).

In addition to the newly formed interactions, we examined how the new binding interface may be further favourable over the wild-type interface. Examination of the difference electron density map of PDB 6M0J reveals that although RBD Q498 can hydrogen bonding with ACE2 Q42 (Fig. 3d), the positive difference electron density adjacent to the side chain is consistent with disorder (i.e. it adopts at least one alternate conformation) (Sup. Fig. 1). Alternate conformations would break the hydrogen bond with ACE2 Q42 suggesting that the contribution of RBD Q498 to ACE2 binding is reduced.

To test whether the H498 imidazole ring structure contributes to the affinity gain, we examined the effect of introducing arginine at position 498, as this would be expected to form hydrogen bonds, but not contribute to any π-interactions. RBD-Q498R was found to have a binding affinity of 7.1nM for ACE2 (Fig 3g), a lower affinity than the 2.0 nM seen with Q498H (Fig. 2c). This shows that the presence of the imidazole ring provides an additional contribution to the affinity gain compared with arginine at this position.

### Effects of Q498H on ACE2 binding of COV-2 RBD variants

The finding that substituting histidine at position 498 in CoV-2 spike protein causes a dramatic increase in binding affinity for ACE2 prompted us to test whether this substitution could enhance binding of other CoV-2 RBD variants. We therefore added the Q498H substitution to two prominent RBD variants, one containing the substitutions L452R and E484Q, which are common to variants B.1.617.1 (Kappa) and B.1.617.3, and one containing RBD substitutions K417N, E484K and N501Y, found in variant B.1.351 (Beta). Addition of Q498H to B.1.617.1/3 (Delta) increased affinity by around three-fold for ACE2, decreasing K_D_ from 7.1nM to 2.5nM (Fig. 4a).

**Figure 4.**
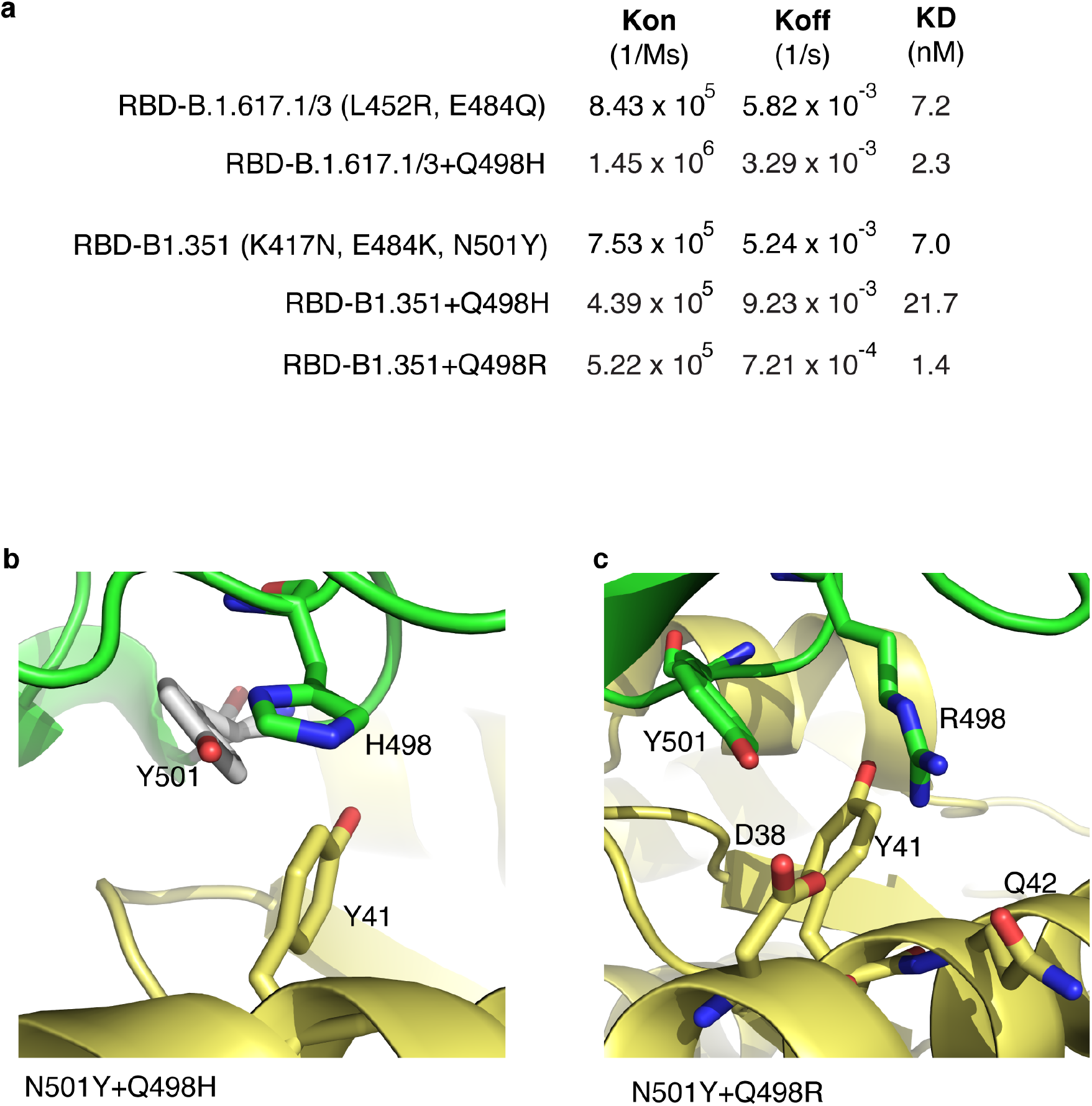
Effects of Q498H and Q498R substitutions on binding of B.1.351 and B.1.617.1/3 variant RBD to HuACE2. **(a)** Biolayer interferometry was performed with the indicated RBD in solution and HuACE2 immobilise on the sensor. Curves were fitted and used to calculate K_on_, K_off_ and K_D_. **(b)** H498 and Y501 in CoV-2 RBD compete for interaction with Y41 in HuACE2. The proximity of the H498 side chain is shown with respect to the side chain of a Y501 inserted into our RBD:ACE2 complex structure. **(c)** R498 and Y501 side chain positions shown along with D38, Y41 and Q42 (PDB accession number 7BH9^16^).

Surprisingly, substitution of Q498H to the B.1.351 variant RBD reduced the affinity of B.1.351 (Fig. 4a). However, H498 and Y501 are in close proximity and the local arrangement required to accommodate the phenolic group of Y501 will be constrained by the imidazole of H498, reducing the advantageous π-π stacking between Y501 and Y41 in B.1.351 as well as perturbing the H498-Y41 interactions (Fig. 4b), consistent with this decreased in affinity. As introduction of a histidine at position 498 clashes with Y501, we tested the effect of an arginine substitution at 498 (Fig. 4a). Addition of Q498R to the N501Y-containing variant increased affinity five-fold, yielding an K_D_ of 1.4nM, similar to that seen with RBD4.8. This increase in affinity is due to a large decrease in off-rate, indicating the Q498R substitution in the N501Y variant acts to stabilize binding to ACE2. Examination of a previously published RBD:ACE2 structure^16^ in which RBD has Q498R and N501Y substitutions, in addition to several other substitutions, suggests rbdY501 makes a π-interaction with aceY41 and rbdR498 forms a hydrogen bond with aceQ42 and a potential salt bridge with aceD38 (Fig. 4c). Therefore, the mechanism by which the combined substitutions of Q498R plus N501Y increases affinity has similarities to that by which the single Q498H substitution increases affinity.

Overall, these data show that the affinity of B.1.617.1/3 variant RBD for ACE2 is enhanced by the addition of the Q498H substitution. In contrast, in a variant which already has an N501Y substitution, the Q498H substitution clashes with Y501 and fails to increase affinity. However, arginine can be accommodated at 498 in a N501Y variant, and this results in a large affinity gain by a bonding mechanism requiring both R498 plus Y501, that shares similarities with that of H498 alone.

### The Q498H/R substitution enables RBD binding to rat ACE2

RBD-Wuhan-Hu-1 is unable to bind rodent ACE2. However, in a recent study to develop a mouse model of CoV-2 infection, SARS CoV-2 was passaged multiple times through mice to derive a variant capable of murine infection^17^. The resultant variant had two substitutions in RBD, Q493K and Q498H, and was capable of binding to mouse ACE2 and causing CoV-2 infection^17^. Interestingly, either of these substitutions alone appears sufficient to allow RBD to bind mouse ACE2. This finding suggests that CoV-2 variants that include a Q498H substitution would be able to bind mouse ACE2 and allow the virus to extend its species range of infectivity. We therefore tested whether the high affinity RBD variant RBD4.8 can bind rodent ACE2. Given the similarity between mouse and rat ACE2 (RaACE2) in the region corresponding to the CoV-2 binding site (Fig. 5b), and the importance of rats as a potential viral reservoir, we focussed on testing binding to rat ACE2.

**Figure 5.**
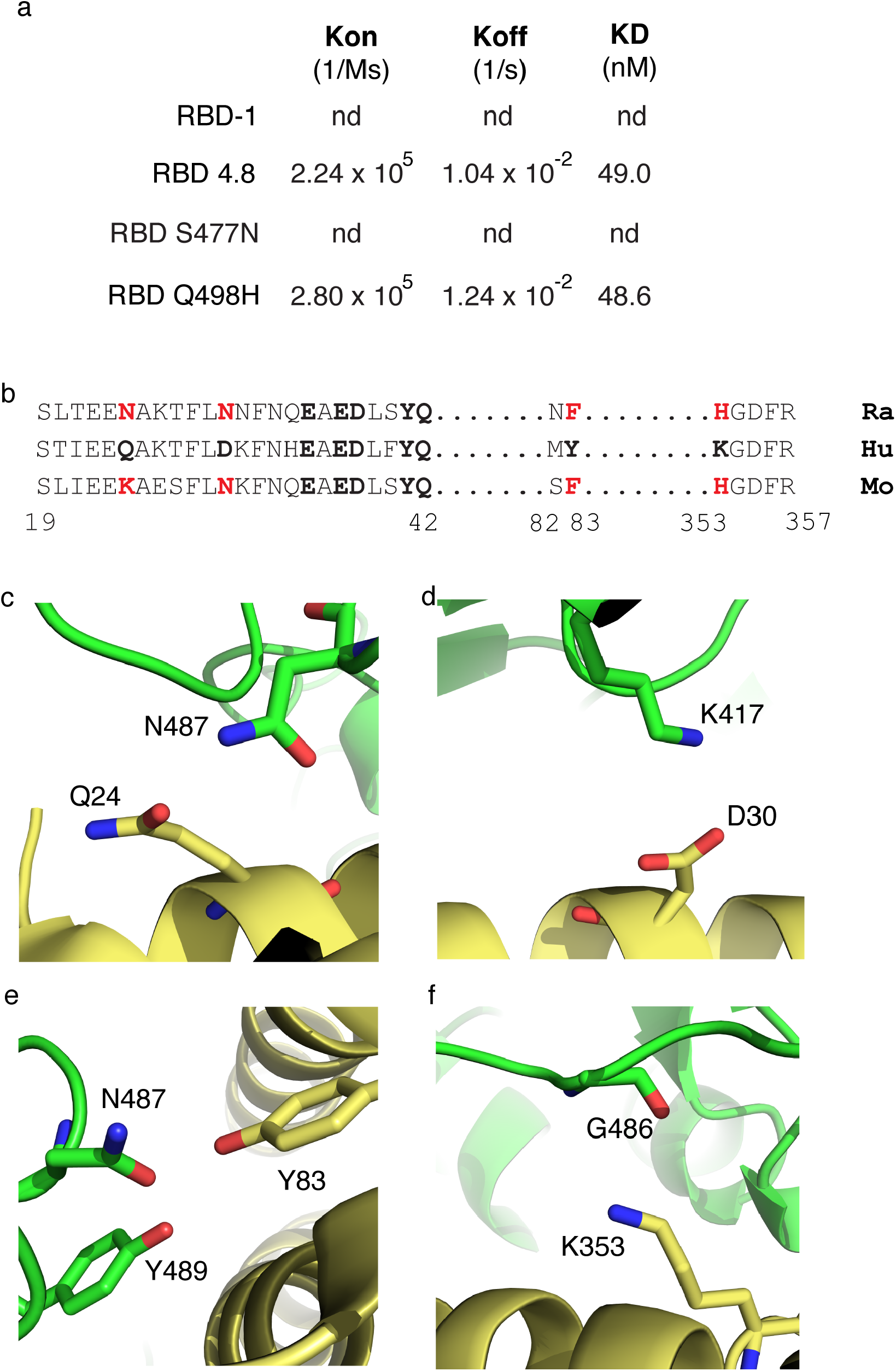
The Q498H substitution enables binding of CoV-2 RBD to rat ACE2. **(a)** Kinetic binding constants for RBD binding to RaACE2 measured using biolayer interferometry. RBD with the Q498H substitution binds rat receptor. **(b)** Rat (RaACE2) and mouse ACE2 (MoACE2) differ from human ACE2 (HuACE2) in key residues contributing to CoV-2 RBD binding. Residues in human ACE2 at the RBD binding interface are aligned with the corresponding residues in rat and mouse ACE2. Residues that form hydrogen bonds or salt bridges with RBD are in bold and those that differ in rodent are in red. (c) Bonding interactions between Q24 and N487 (d) D30 and K417 (e) Y83 and N487 plus Y489, and (f) K353 and G496, in HuACE2 (yellow) and RBD (green).

Like the situation with mouse ACE2, we could not detect any binding between RBD1 and RaACE2 (Fig. 5a). However, we found that RBD4.8 was clearly capable of binding RaACE2, with a K_D_ of 49nM, which is comparable to the affinity of Wuhan-Hu-1 RBD for human ACE2 (Fig. 2c). RBD with the S477N substitution alone was unable to bind the rodent receptor, whereas RBD with just the Q498H substitution bound with an affinity similar to RBD4.8 (Fig. 5a).

### Mechanisms of binding to rodent ACE2

Binding of RBD to HuACE2 involves three key regions of ACE2, the alpha helix between residues 19 and 42, residues 82 and 83 and residues 353 to 357^18-20^. Alignment of the ACE2 sequences encompassing these regions for human and rat reveals differences between the species (Fig. 5b). Four residue positions identified as being involved in salt bridge or H bonding between HuACE2 and CoV-2 RBD differ in the rat sequence, Q24, D30, Y83 and K353 (Fig. 5b). Of these, Q24 in HuACE2 hydrogen bonds with rbdN487 (Fig. 5c), and this hydrogen bond will also likely be formed by rat ACE2 where position 24 is asparagine. In contrast, aceD30 forms a salt bridge with rbdK417 (Fig. 5d) and this would not occur with the corresponding N30 in rat ACE2. Y83 forms two hydrogen bonds with N487 and Y489 (Fig. 5e) and these would also be abolished in rat ACE2 where corresponding residue at position 83 is replaced by phenylalanine. K353 in HuACE2 hydrogen bonds with G496, however in rat 353 position is occupied by histidine and the shorter reach and lower mobility of the histidine side chain would limit the ability of H353 to form this hydrogen bond. Overall, this loss of a salt bridge and at least two H bonds would substantially decrease binding affinity resulting in the apparent lack of binding between CoV-2 RBD and RaACE2.

Examination of mouse ACE2 over the key residue positions involved in HuACE2:RBD binding shows four residue positions involved in HuACE2:RBD binding differ in the mouse sequence (Fig. 5b). Of these, three are the same as the residues found in rat ACE2, namely N30, F83 and H353 and would, as in rat, result in loss of a salt bridge and at least two hydrogen bonds. In addition, position 24 is a lysine in mouse ACE2, rather than the glutamine or asparagine found in human and rat, and would result in loss of the hydrogen bond with N487. Therefore, like rat, loss of key hydrogen bonding and salt bridge interactions would result in a lower in binding energy and explain the inability of RBD1 to bind detectably to mouse ACE2.

The ability of Q498H RBD to bind RaACE2 indicates this substituted RBD can form additional inter-molecular bonds, compared with RBD1. As shown in Figure 5b, two of the three residues we found in HuACE2 to be bonding with H498 are conserved in RaACE2, namely D38 and Y41. Thus, H498 could form a π-interaction with aceY41 and a hydrogen bond with aceD38, as in HuACE2, providing additional binding energy not available to RBD1. D38 and Y41 are also conserved in mouse ACE2 (Fig. 5b), potentially allowing H498 to, similarly, form additional bonds. Binding of mutant RBD with a Q498H substitution to rat, as well as mouse^17^, ACE2, therefore, likely reflects an enhancement of affinity, due to H498 bonding with Y41 and D38, bringing binding of RBD to rodent ACE2 up to a detectable level, and thus enabling CoV-2 variants with Q498H substitutions to infect rodents^17^.

### Q498H and Q498R + N501Y enable binding of RBD to rat ACE2

We were interested to test whether B.1.617.1/3 and B.1.351 variant RBDs can bind RaACE2, and whether this is affected by acquisition of an additional basic substitution at Q498. As shown in Table 1 we found no detectable binding of the B.1.617.1/3 variant RBD to RaACE2. However, addition of the Q498H substitution results in an increase in affinity for RaACE2 resulting in detectable binding of this variant, with a K_D_ of 29.7nM. RBD variant B1.351 did not bind detectably to RaACE2 (Table 1). As substitution Q498H was found to clash with N501Y (see above), we tested the effect of addition of Q498R to the B1.351 RBD. This additional substitution increased binding to detectable levels, resulting in a K_D_ of 36nM for RaACE2, whereas Q498R alone did not produce detectable binding to RaACE2 (Table 1).

**Table 1.** Effects of Gln498His and Gln498Arg substitutions on binding of B.1.617.1/3 and B.1.351 variant RBD to RaACE2. Biolayer interferometry was performed with the indicated RBD in solution and RaACE2 immobilised on the sensor, nd indicates signal not detectable.

| **RBD Mutations** | **K<sub>on</sub> (1/Ms)** | **K<sub>off</sub> (1/s)** | **K<sub>D</sub> (nM)** |
| --- | --- | --- | --- |
| B.1.617.1/3 (L452R, E484Q) | nd | nd | nd |
| B.1.617.1/3 + Q498H | 2.60 x 10<sup>5</sup> | 7.60 x 10<sup>-3</sup> | 29.7 |
| B.1.351 (K417N, E484K, N501Y) | nd | nd | nd |
| B.1.351 + Q498R | 3.20 x 10<sup>5</sup> | 1.14 x 10<sup>-2</sup> | 36.4 |
| Q498R | nd | nd | nd |

## Discussion

In this study we have used the DT40 directed evolution system to identify a readily accessible double mutant form of CoV-2 spike protein RBD (S477N/Q498H) with a large enhancement in binding affinity for HuACE2, which is therefore likely to increase transmissibility of CoV-2 in humans. Furthermore, we show that the Q498H substitution, and the related Q498R substitution plus N501Y, enables binding of RBD to rat ACE2. We provide a molecular mechanism for the increased affinity of the double mutant and also show that Q498H, or the related Q498R substitution, confers large affinity gains on other CoV-2 RBD variants. The Cryo-EM structure of the RBD4.8:ACE2 complex revealed the newly formed interactions of a histidine at position 498, including π-stacking and hydrogen bonding to neighbouring ACE residues, strengthens the previous transient interactions between Q498 and ACE2. In addition, a new hydrogen bond between N477 and ACE2 at the other end of the RBD binding interface contributes to the almost 20-fold binding increase. Nonetheless, the increase in binding is driven mainly by the Q498H. Indeed, Histidine at position 498 results in an affinity gain greater than that reported for most other single mutations in RBD. Consistent with this idea, Starr *et al*^9^, using deep mutational scanning, show Q498H to be the most enriched substitution in their highest binding RBD pool.

We hypothesised that this substitution would increase the binding affinity of other CoV-2 variant RBDs, and indeed gain of Q498H resulted in a clear enhancement of binding in B.1.617.1/3 variant RBD. Thus, acquisition of Q498H would be expected to increase binding and transmissibility of these viral variants. In contrast, addition of Q498H to B.1.351 variant RBD decreases affinity, raising the K_D_ for ACE2 from 7nM to 21.7nM (Table 1). The reason for this is evident from structural analysis, which reveals close proximity of H498 to the Y501 substitution in B.1.351, causing a clash of the side chain rings for binding to ACE2 residues.

However, we find that a variant RBD with tyrosine at 501 can accommodate an arginine at 498, enhancing affinity by more than six-fold in the N501Y RBD we tested. Interestingly, Q498R results in only a small increase in affinity in the absence of a tyrosine at 501 in RBD. This suggests Q498R substitutions in CoV-2 would be more likely in lineages in which N501Y is already fixed. Indeed, in a recent *in vitro* evolution study, the Q498R substitution was only evident after early fixation of N501Y and high selection pressure for affinity^16^.

CoV-2, therefore, can gain a substantial affinity boost by two different routes both involving position 498, and substitution preferences between Q498H and Q498R are determined by whether N501Y is already present in the spike protein. CoV-2 RBD lineages with an N501Y substitution would be unlikely to gain Q498H without losing fitness, whereas such variants could gain Q498R leading to increased affinity. Significantly more than 85% of reported CoV-2 variants containing the Q498R substitution also have the N501Y substitution, and the vast majority of these are the recently described B.1.1529 variant (Omicron), that also contains S477N and a range of other substitutions^21^. This variant is expanding rapidly and based on our findings, we anticipate that Q498R plus N501Y, as well as S477N, make significant contributions to the binding ability, human-to-human and potential cross-species transmissibility of Omicron. Our findings also mean that CoV-2 lineages without N501Y could gain a corresponding substantial affinity increase by acquiring the single Q498H substitution, potentially enabling parallel viral lineages with either Q498H or Q498R plus N501Y as affinity contributors.

Ordinarily, Wuhan-Hu-1 CoV-2 RBD is unable to bind detectably to rodent ACE2 and here we show why this is the case. However, we find that substitutions at Q498 opens up RBD binding to rat ACE2, our analysis pinpointing the D38, Y41 ACE2 residue dyad as a key site exploited by the virus for the enabling affinity gain. Addition of variant-specific Q498 substitutions also confer rat ACE2 binding on CoV-2 RBD variants. In our experiments, gaining a Q498H, or Q498R plus N501Y, resulted in binding of RBD variants that were otherwise incapable of detectable binding to rat ACE2. These mutations, already found in Omicron and other emerging CoV-2 variants, thus have the potential to both increase human transmissibility and extend these variants into rodent populations. Acquisition of mutations that enable CoV-2 to bind rat ACE2 is a cause of concern, as this has the potential to facilitate transmission of the virus to a species that is widespread and lives close to humans. Although, whether interactions between humans and rats would be sufficient to allow cross-species transmission is not yet known. Further evolution in such a reservoir carries the additional potential risk of a spill back to humans of novel variants with further detrimental phenotypes. Worryingly, CoV-2 variants with Q498H, and Q498R plus N501Y, have already been detected in wastewater^11^, a potential transmission route to rat.

## Methods

### Materials

DNA encoding the N-terminal secretory leader sequence from CD5 (residues 1-24) upstream of either HuACE2 (residues 19-615) or RaACE2 (residues 19-165), followed by a short linker FLAG-epitope tag and C-terminal Histidine_6_ was synthesised by GeneArt (Invitrogen). To generate a FLAG-free HuACE2 fusion, the FLAG-epitope tag in the HuACE2 construct was replaced with a HA tag. All other reagents were as described previously^11,22^.

### Cell surface display

The cell surface display DT40 system that we previously described^10^ was used. cDNA encoding a fusion protein comprising of an N-terminal CD5 secretory leader sequence followed by the Wuhan-Hu-1 CoV-2 RBD (residues 319-541), linker region and FLAG epitope tag together with a C-terminal transmembrane domain and short intracellular domain fragment of platelet-derived growth factor receptor-β was synthesised (GeneArt Gene Synthesis) and inserted into the pHypermut2 vector^23^ This was transfected into DT40 cells by electroporation and stable transfectants derived by growth in puromycin. Clonal DT40 lines in which the RBD construct was integrated into the rearranged Ig locus were identified by PCR, and surface expression of the fusion construct confirmed by anti-FLAG immunostaining as previously described^10^. Cells were cultured in RPMI containing 7% (v/v) fetal bovine serum (FBS) plus 3% (v/v) chicken serum at 37°C and 5% CO_2_.

For ACE2 binding to DT40 cells, approximately 40 million cells were washed and incubated with 0.1nM Histidine_6_-tagged HuACE2 in PBS with 10% (v/v) FBS at room temperature for 30 mins. Cells were washed and incubated with anti-FLAG-APC and anti-Histidine_6_-PE antibodies on ice for 20 mins, followed by washing. Cells with bound ACE2 were selected by fluorescence activated cell sorting on a FACS Aria Fusion (Becton Dickinson) with the sort windows indicated in Results. Sorted cells were resuspended and grown in DT40 culture medium. For sequencing, genomic DNA from an aliquot of the sorted cell population was recovered using PureGene DNA isolation kit (Qiagen) and RBD amplified by PCR. Purified PCR products were directly sequenced. In addition, purified PCR products were inserted into pcDNA3.1, transformed into *E. coli* and sequencing was performed on randomly picked colonies.

### Site-directed mutagenesis and soluble Fc-fusion proteins

Site directed mutagenesis was performed using the QuikChange protocol (Agilent Technologies) and constructs were sequenced to confirm mutations. cDNA encoding Fc-fusion proteins were constructed by ligating the appropriate RBD or ACE2 nucleotide sequence upstream of a GS4 linker, fragment of human Fc immunoglobulin domain and C-terminal Histidine_6_ tag.

### Expression and purification of soluble proteins

HEK 293 cells were transfected with mammalian expression vectors encoding the relevant protein using polyethylenimine and cells incubated for approximately 3 days to allow accumulation of the secreted protein in the medium. Harvested media was centrifuged and filtered and H_6_-tagged proteins recovered by nickel chromatography. After washing of columns, proteins were eluted with imidazole and Zeba columns used for buffer exchange into Tris-buffered saline (TBS; 25mM Tris, 150mM NaCl) containing 10% glycerol (v/v). Protein purity was assessed by SDS polyacrylamide gel electrophoresis and Coomassie staining. Bradford assays were performed to determine protein concentrations and proteins were stored at 4°C before use.

### Biolayer interferometry

Binding analysis was performed by biolayer interferometry using an Octet RED instrument. Assay buffer comprised of tris-buffered saline with 0.05% (v/v) Tween-20 and 1mg/ml BSA. AHC biosensors were hydrated and then coated with ACE2-Fc by dipping sensors into assay buffer containing 5μg/ml ACE2 for 150 sec. After washing, association was measured by immersion of coated sensors in assay buffer containing the concentrations of soluble RBD monomers indicated in Results, followed by return of the sensors to buffer to measure dissociation. Data was analysed using the Octet Data Analysis Software with a 1:1 binding model.

### Flow cytometry

Vero-E6 cells were cultured in DMEM with 10% (v/v) FBS. For cellular binding analysis cells were collected by centrifugation, washed and incubated for 15 min at 37°C with 5μg/ml of the appropriate HA-tagged RBD in PBS with 10% (v/v) FBS. After washing bound RBD was detected by incubating cells for 10 min at room temperature with anti-HA conjugated to PE. Cells were then washed and mean fluorescence of stained cell populations determined on a FACS CantoII flow cytometer (Becton Dickinson).

### Complex preparation

The complex of ACE2 and mutant RBD used for cryo-EM was formed at room temperature for 30 min, concentrated using an Amicon Ultracel 10K Centrifugal Filter, cleared through a Millipore 0.22 micron Durapore centrifugal filter and applied to a Superdex-200™ (10/300 GL) column (GE Healthcare) which was pre-equilibrated and then run in gel filtration buffer (TBS). Fractions were collected every 250 μl, monitored by *A*_230_ and analysed by SDS-PAGE (NuPAGE™ 4-12% Bis-Tris). Fractions containing the complex were selected and protein concentration determined using the extinction coefficient for the 1:1 complex at *A*_280_. The fraction containing the highest concentration of complex was then used for subsequent analysis by Cryo-EM.

### Cryo-electron microscopy

Grids were prepared on either unsupported holey grids or holey grids overlaid with graphene oxide. Samples derived from size exclusion chromatography were used without further concentration at 0.2mg/ml on holey grids or diluted to 0.05 mg/ml for graphene oxide grids. For the former, grids were glow-discharged for 60 sec at 35 mA on a Quorum GloQube. Graphene oxide grids were prepared as described before^22^ and were glow-discharged at 40 mA for 180 sec prior to graphene oxide application. In each case 3μl of the complex was applied to grids (Quantifoil R1.2/1.3 300 mesh Au) and plunge frozen using a Thermo Fisher Scientific Vitrobot MKIV. A wait time of 30 sec was applied for graphene oxide grids to allow particles to adhere to the support film.

Data were collected on a Thermo Fisher Scientific Titan Krios G3 operating at 300 KeV and equipped with a Gatan BioQuantum energy filter with a slit width of 20 eV and a K3 direct electron detector. Movies were recorded at a nominal magnification of 105Kx in Counting Bin1 mode using aberration free image shift (AFIS). A total dose of 50e-/Å^2^ fractionated over 50 frames and a defocus range of -0.7μm to -2.7μm in 0.3 intervals were used. We proceeded to image the complex by Cryo-EM but initial attempts using holey grids resulted in strong preferential orientation and poor-quality maps (Sup. Fig. 2a). When the complex was imaged on grids overlaid with graphene oxide we also observed a strong orientational bias in a different direction (Sup. Fig.2b). Combining the two datasets however resulted in a good quality 3.2Å map (Fig.3a).

### Image processing and Model Building

All data processing was performed using Relion 3^24^. Briefly, movies were corrected for motion using MotionCor 2.1.4^25^ and the contrast transfer function parameters were estimated using GCTF 1.18^26^. Particles were picked automatically using Topaz^27^ initially with the supplied model but later on using a trained model based on the data. Initial processing of the unsupported dataset indicated severe preferential orientation (Sup. Fig. 2) and despite efforts to computationally balance the dataset a good 3D map could not be obtained. A second dataset on graphene oxide also indicated strong preferential orientation in a different direction so the two datasets, collected under identical optics conditions, were combined and processed together. After several rounds of cleanup using 2D classification, the data were subjected to 3D classification and the best subset was chosen for further refinement. The data was further improved by refining CTF parameters and aberrations and particle polishing. A final map was obtained at a global resolution of 3.2Å using the Gold Standard FSC 0.143 criterion.

Initial rigid-body docking of the crystal structure (PDB ID 6M0J) was performed using UCSF Chimera 1.15^28^ and further model building was performed in Coot 0.9.6^29^. After manual rebuilding real-space refinement of the coordinates was performed using Phenix 1.19.2^30^. All figures were created sing Chimera X 1.3rc^15^.

The Cryo-EM maps and coordinates have been deposited to the EMDB (EMD-XXXX) and PDB (PDB ID XXXX) respectively.

## Acknowledgements

This work was funded by the Medical Research Council (MC_PC_19043). NB and NPJB are supported by the British Heart Foundation (PG/19/27/34305). PCEM is a Royal Society Wolfson Fellow (RSWF\R3\183003). Work in the JES group is supported by a core grant to the LMB from the MRC (U105178808). We acknowledge the Midlands Regional Cryo-EM Facility at the Leicester Institute of Structural and Chemical Biology (LISCB), major funding from MRC (MC_PC_17136).

## Author contributions

NPJB and JES conceived and supervised the project: NPJB, JES, NB, CGS, PCEM, JKB and JWRS provided conceptual input: NB, NPJB and AEB performed experimental work; CGS prepared Cryo-EM samples, performed data collection and processing: CGS, PCEM, JWRS performed Cryo-EM model building and structural interpretations: all authors contributed to manuscript writing and revision.

