## Supplementary Figs for "*In vitro* evolution predicts emerging CoV-2 mutations with high affinity for ACE2 and cross-species binding"

**Supplementary Figures**

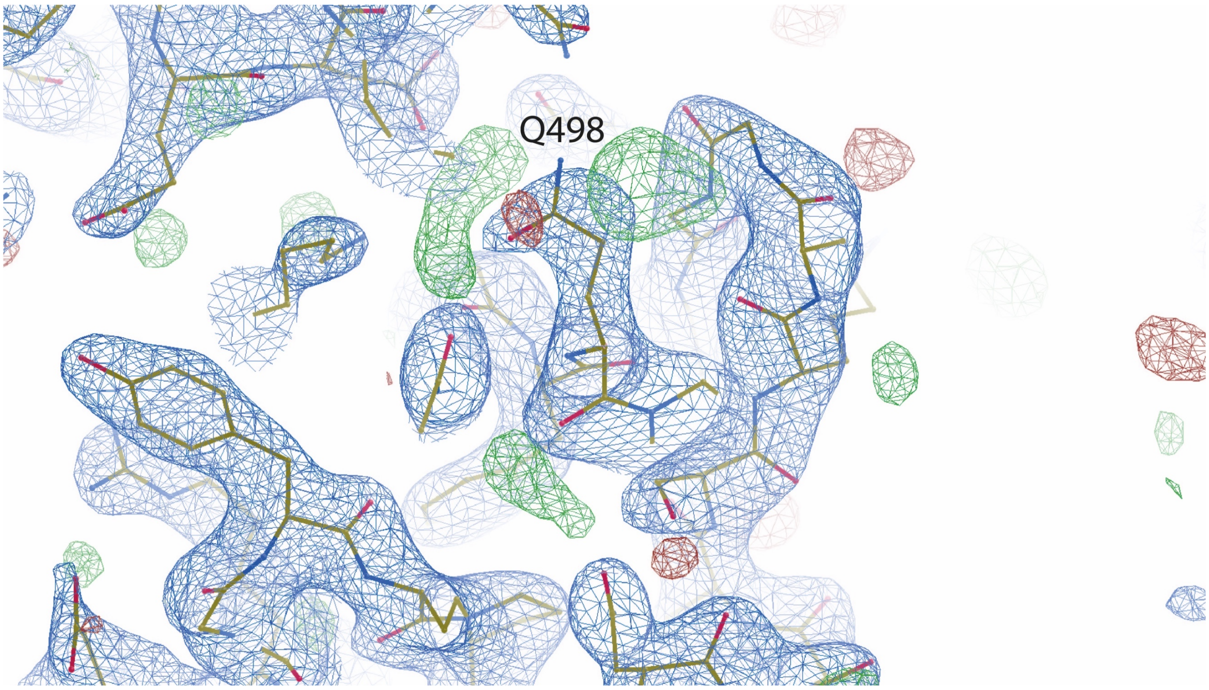


**Sup. Fig. 1. Multiple conformations of Q498 are suggested in WT crystal structure.** The 2Fo-Fc electron density map (in blue, contoured at 1.5σ) and Fo-Fc difference map (contoured at -3σ (red) and green (+3σ)) are shown for the WT crystal structure of ACE2-RBD (PDB ID 6M0J[1]). The positive (green) density adjacent to the side chain of Q498 suggests this can exist in different conformations, thus weakening the interaction with neighbouring residues.

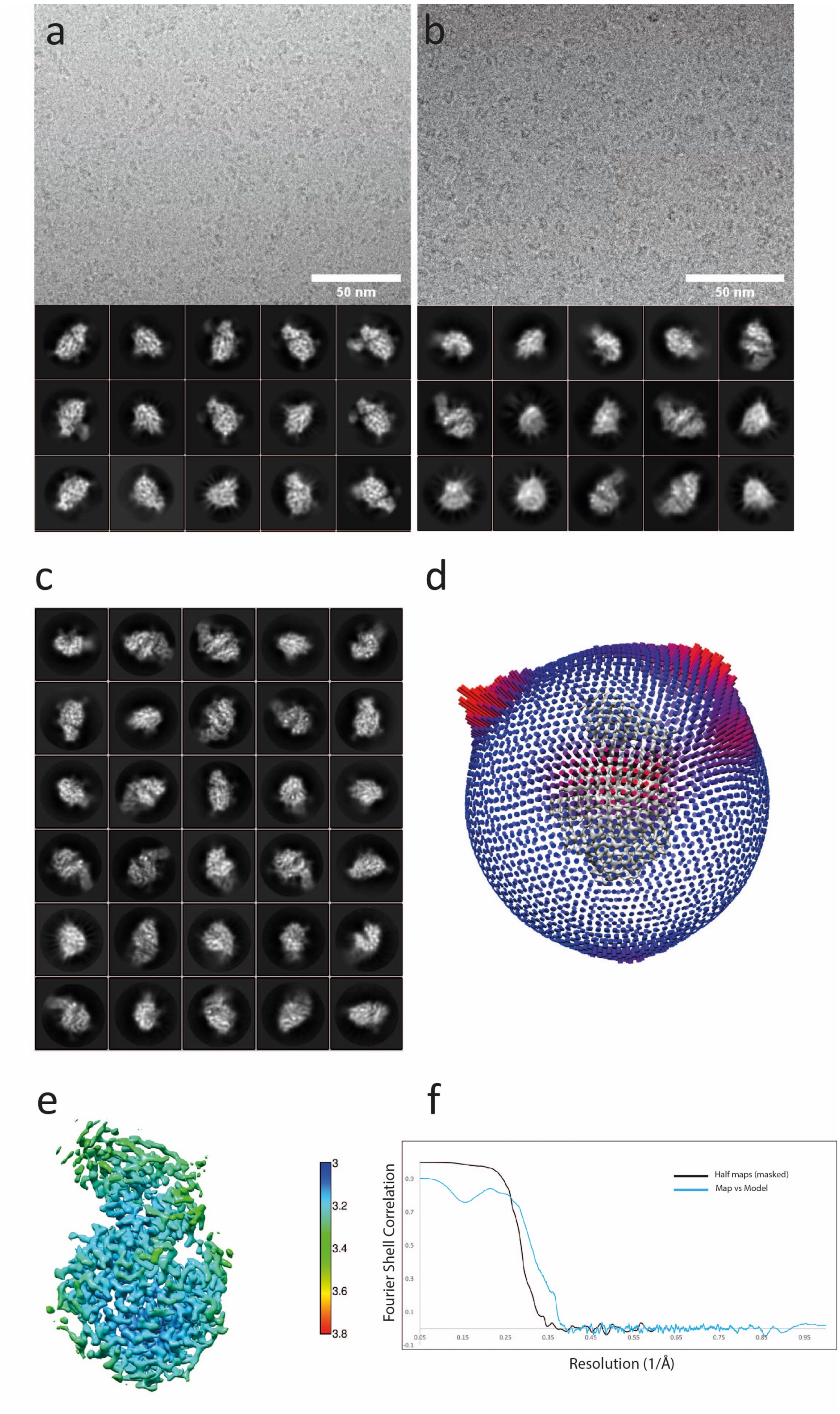


**Sup. Fig. 2 Cryo-EM of the HuACE2-RBD4.8 complex.** Representative micrograph and 2D classes of the complex on either unsupported holey grids (a) or graphene oxide coated holey grids (b). The combined datasets result in the 2D classes shown in (c) and the angular distribution shown in (d) with an overall uniform angular distribution despite some persistent angular bias. The local resolution map is shown in (d) and the FSC of the two half maps and map vs model are shown in (f).
